# Efficient inference of single cell expression profiles with overlapping pooling and compressed sensing

**DOI:** 10.1101/338319

**Authors:** Xingzhao Wen, Weiqiang Xu, Xiao Sun, Jing Tu, Zuhong Lu

**Author notes:** Correspondence (J.T.), (Z.L.).

## Abstract

Plate-based single cell RNA-Seq (scRNA-seq) methods can detect a comprehensive profile for gene expression but suffers from high library cost of each single cell. Although cost can be reduced significantly by massively parallel scRNA-seq techniques, these approaches lose sensitivity for gene detection. Inspired by group testing and compressed sensing, here, we designed a computational framework to close the gap between sensitivity and library cost. In our framework, single cells were overlapped assigned into plenty of pools. Expression profile of each pool was then obtained by using plate-based sequence approach. The expression profile of all single cells was recovered based on the pool expression and the overlapped pooling design. The inferred expression profile showed highly consistency with the original data in both accuracy and cell types identification. A parallel computing scheme was designed to boost speed when processing the enormous single cells, and elastic net regression was combined with compressed sensing to auto-adapt for both sparsely and densely expressed genes.

## INTRODUCTION

Single cells, as the basic components of life, are a new window to understand individual differences among cells (Janiszewska et al., 2015; Raj et al., 2008). With the development of advanced technologies to capture single cells quickly and accurately (Perfetto et al., 2004; Spitzer and Nolan, 2016), scientists can narrow down their view from bulk sequencing of thousands of cells, which averages out cellular difference, to the subtle changes between individual cells (Zeisel et al., 2015). The elaborately atlas of single cell, has shed light on multiple biological questions like revealing new cell types in cancers (Chung et al., 2017; Grün et al., 2015), investigating the dynamics of developmental processes (Li et al., 2017), linkage and developmental trajectory of immune cells in cancer (Zheng et al., 2017) and identification of gene regulatory mechanisms (Datlinger et al., 2017).

The two most popular methods for single-cell sequencing are plate-based protocols and microdroplet-based methods. Plate-based protocols like SMART-Seq2 (Picelli et al., 2013; Picelli et al., 2014; Tang et al., 2009) have higher sensitivity in gene detection by costly constructing sequencing library for each cell independently. Correspondingly, microdroplet-based methods like Drop-seq (Klein et al., 2015; Macosko et al., 2015) are more efficient in sequencing by building one barcoded library for massive cells to analyze large amount of cells in parallel with low cost. Although this approach is widely used, these droplet-based approaches lose sensitivity when compared with the plate-based methods. For each cell, only about a half of genes can be detected (Ziegenhain et al., 2017). And the 3’ sequencing method cannot capture the full length of RNA which prevents deep downstream analysis (Baran-Gale et al., 2017). In short, the manual library construction methods are superior in sensitivity but suffer from the high cost, which is usually a hundred times more expensive than microdrop-based methods for each cell (Baran-Gale et al., 2017). Therefore, an efficient approach is desirable to be developed to close the gap between cost and efficiency of gene detection.

In this paper, we proposed an improved technique to cut down the budget of library construction while maintained the same sensitivity to capture comprehensive gene expression profiles for scRNA-seq. The basic idea behind our method is to group different cells into overlapped pools, where the number of pools is usually much less than the number of cells. We only construct library for each pool and get composite sequencing results. Certain algorithms are applied to recover expression level for each single cell.

Except some house-keeping genes which routinely expressed cross all kinds of cells, single cell RNA-seq expression profile (SCEP) is usually sparse for most genes due to a lot of drop out events, which is also called zero-inflated (Pierson and Yau, 2015; Vallejos et al., 2017). This intrinsic feature of scRNA-seq data suggests that they are compressible. Our algorithm to recover the expression profile is based on the idea of group testing and compressed sensing methods. The objective of quantitative group testing was first raised to identify counterfeit coin by group test weights, where the weights are analogue to gene expression level (Grossman and Shapiro, 1988). Basic Group testing method has also been applied in many biological problems like rare variant detection (Erlich et al., 2009) and haplotype assembly (Li et al., 2016). Compressed sensing (CS) was first designed to sample and recover the sparse signals at a much lower rate rather than Shannon/Nyquist sampling frequency (Candes et al., 2006). By random sampling, CS method can overcome two handy occasions: where high sampling rate is not practical in some situations (Rani et al., 2018) and the cost of each sample is pretty high. Here we interpret each pool as a measurement to detect our sparse expression profile, whilst use limited sampling rate (less library) to recover the expression table with much lower cost.

Our aim can be divided into two levels. The first one is to approach the approximately same accurate SCEP as the manual library construction method with less libraries. The second is step back a little bit to examine whether we can maintain most of the original data structure with a compromise of losing some accuracy but reduce more library usage (Figure 1).

**Figure 1.**
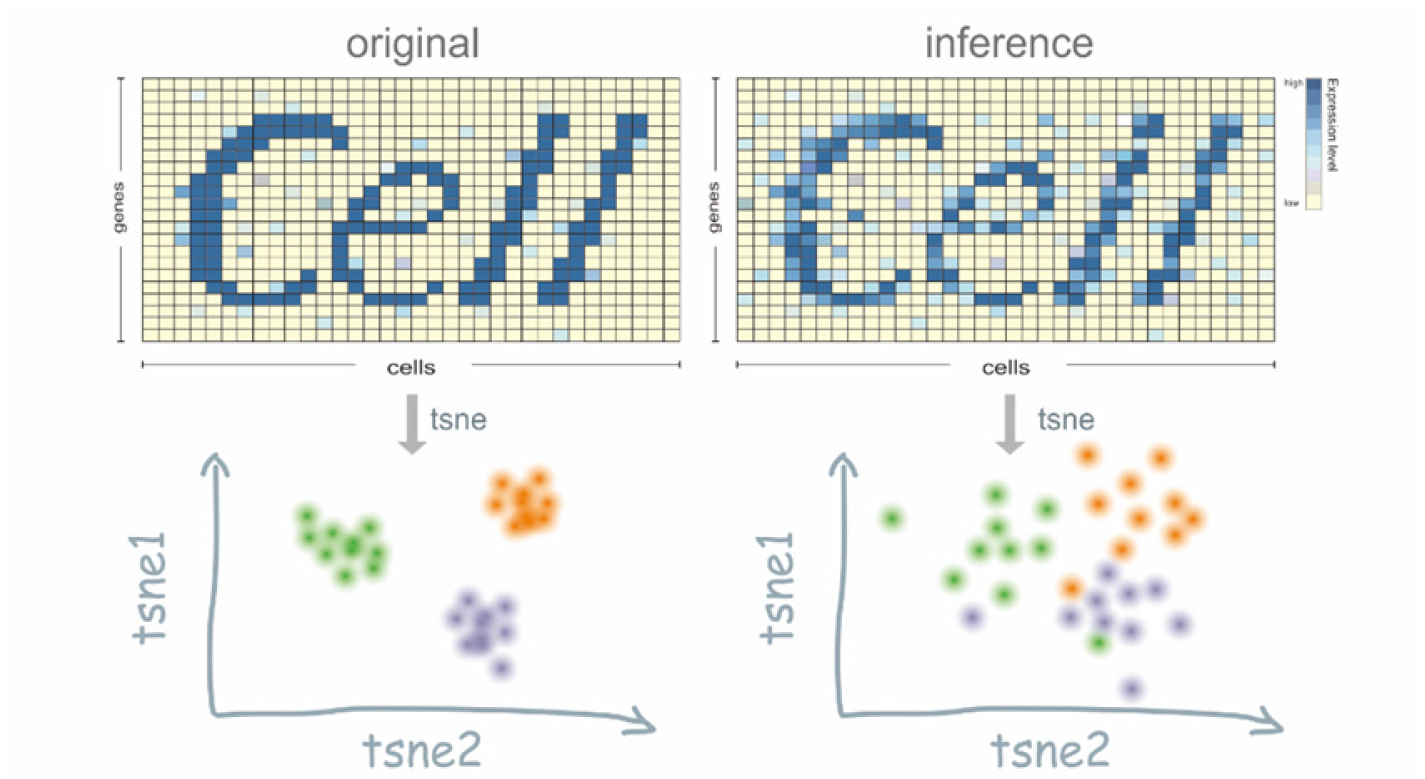
Objection overview illustration. Single cell expression profile is usually sparse at the gene level (row of profile), whereas inference profile is dispersive due to group partition and recovery algorithms. t-SNE (Van Der Maaten and Hinton, 2008) is used for dimension reduction visualization.

A comprehensive computational framework was proposed here to impute SCEP with composite measurements. We first implemented compressed sensing algorithm alone on a plate-based 64 cells SCEP dataset (Li et al., 2016) to show the ability of recovering the data structure by our method. Then, we extended to a much larger dataset containing 5063 single-cells (Zheng et al., 2017) under a parallel computing paradigm by segmenting all cells into sub block, and at the same time evaluated both the accuracy and the maintenance of principle data structure. To cope with dense genes, which usually expressed in various kinds of cells, we proposed a possible solution using self-adapted regression method. Finally, for various considerations, we provided a more accurate model but with a higher demand of experimental labor. The three frameworks were demonstrated to allow users to achieve different resolution of the original expression profile by using different number of pools, while reduce the library construction cost as a whole.

## RESULTS

### From Grouping Idea to Solving Linear Equation: Model Design

To find a systematic approach to achieve our goal of reducing library usage, we learned the idea of group testing method, divided all cells into overlapped groups. Here overlapped means one single cell can appear in different pools, and one pool contains a group of cells. This overlapping fashion utilize the same cell as an information bridge and can reduce the tests by cross information. The sparsity of SCEP which suggests it is compressible. Compressed sensing is a widely used method for sparse signal recovery and sampling. We combined these two ideas to form our model (Figure 2).

**Figure 2.**
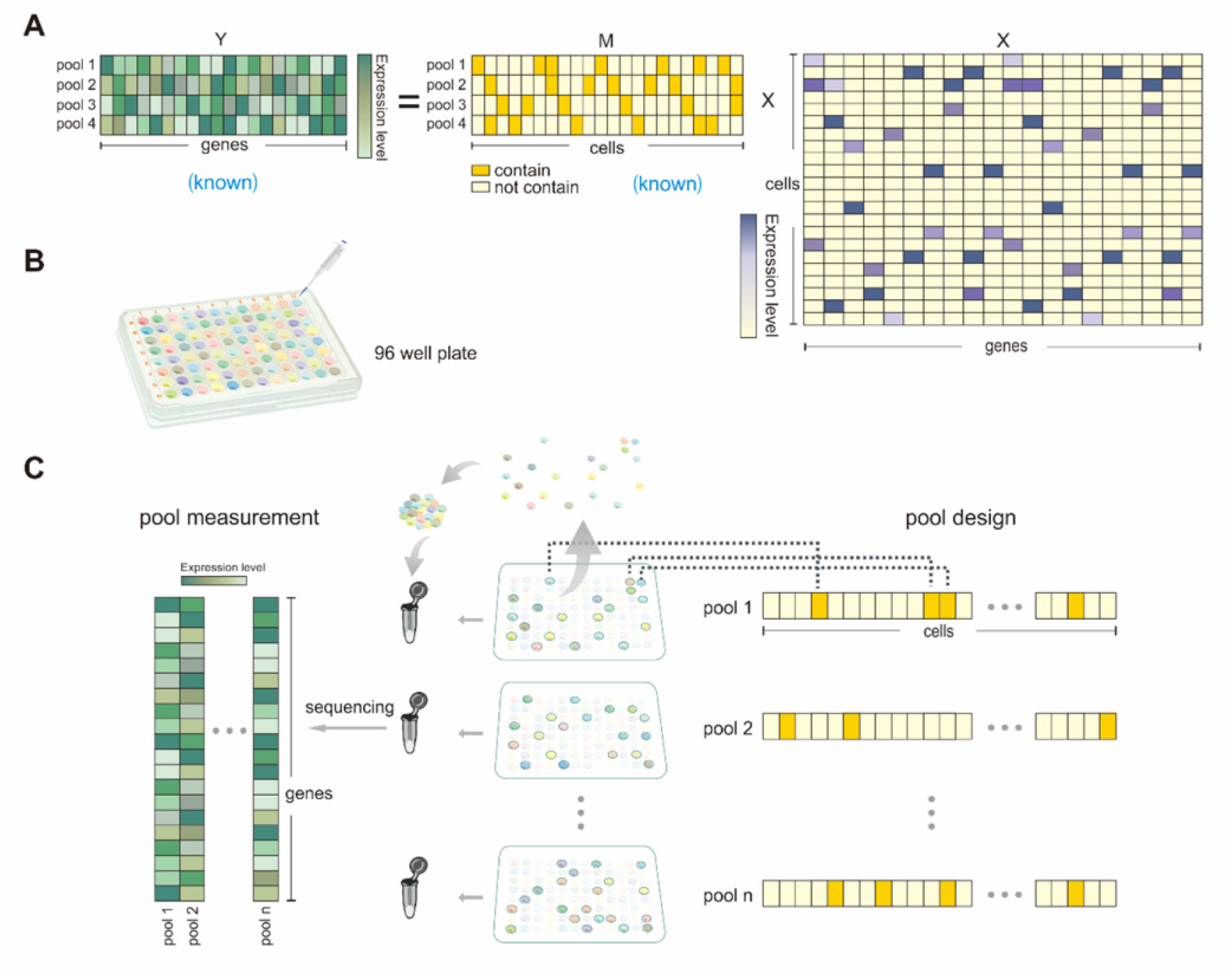
From grouping idea to linear equation. (A) Illustration of pool strategy for single cell expression profile inference. The M matrix is pool-cell matrix, which is overlapped. In this example, 22 cells are designated into 4 pools. M_11_ is orange means pool #1 contains cell #1. Y is pool-gene matrix, which is the expression level for each pool after sequencing. Different colors represent different levels. X is cell-gene matrix, that we want to infer from Y and M. It is usually sparse among most genes. (B) Illustration of pooling strategy, same amount of lysate of each cell is extracted into pools. (C) In the middle, different masks on the 96 well plate represent different cell group in each pool. Each plate is an 8*12, on the right is a long one-dimensional vector with length of 96. It contains value zero (white) and one (orange) indicating how to choose cells for each pool. On the left, a grouping-based sequencing result for each pool.

Pooling (or grouping) is the central idea of our method, and the way we divide different cells into different pools is represented with a pool design matrix 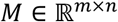. In the basic scenario, suppose *M* = {*m_ij_*} is a binary *m* × *n* matrix, where *m* indicates the number of pools which equals the number of library we will use, and *n* represents the number of cells. *m_ij_* is binary and *m_ij_* = 1 if and only if *i^th^* pool contains *j^th^* cell (Figure 2A).

Pools are designed according to the *M* matrix first, and then sequenced using NGS technology. The result is represented by the 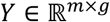 matrix, which is the expression profile for pools. Suppose the whole SCEP is 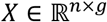, then *Y_:j_* = *M* × *X_:j_* can be easily interpreted as: we get *Y_:j_* (a compressed format of gene *j*’s expression profile) by using a combination of detectors *M* to overserve the SCEP *X_:j_* (original format of gene *j*’s expression profile). This question can be transformed into a matrix form (Figure 2A):

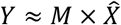

Our goal is to solve this equation effectively in high accuracy with a properly designed *M*.

Mathematically, this ill-conditioned linear equation (*m* is usually much less than *n*, which meets our needs to use less library) has infinite answers. However, compressed sensing can help to solve this equation when *X* is sparse. A well-defined *M* which satisfies some requirements can recover an accurate *X* with confidence. The requirements on *M* can be relaxed if we just want a silhouette of the SCEP.

Experimentally, since one single cell will be sequenced many times, people can lyse single cells at the beginning and use aliquots of cell lysate to represent the exact cells. Each pool contains several cells, and each cell appears in different pools (Figure 2B, Figure 2C). Suppose we want to sequence 96 unique cells using a pool strategy, we would first lyse each cell into different wells of a 96 well plate (Figure 2B), and then adopt a different combination of different cells for each pool. The same amount of cell lysate is extracted into a test tube, therefore one library is constructed for one tube and then sequenced a whole tube at a time.

### Toy Example Using Compressed Sensing Maintained Core Structure

We first applied compressed sensing method alone on a 64 single-cell dataset with 17,683 genes (Li et al., 2016) to evaluate the recovery ability. This dataset contains 64 cells from human pancreas and were sequenced based on plate-based Smart-Seq2 protocol (Picelli et al., 2014).The dataset contains 7 defined pancreatic cell types and 1 undefined group. We recovered the SCEP and visualized with t-SNE (Hinton, 2008; Krijthe, 2015) with given labels from original dataset (Li et al., 2016) to verify the performance.

Two measurements were used to evaluate the performance: (1) Pearson correlation for each gene between the original SCEP and the inferred one. (2) A low-dimensional embedding representation. As expected, the overall correlation calculated based on all genes was increased with the increment of both pool number and cell number of each pool (Figure 3A). However, when taking two variables as a whole, the pool numbers played a more important role, as decreasing the pool numbers from 20 to 15 led to a dramatic fall in correlation from 0.98 to 0.83 (32 cells in one pool). This decreasing trend is also non-linear. In general, using 30 pools can capture most information which is highly coherent with original data, which is visualized by low dimensional embedding (Figure 3B). Delta, beta, acinar and alpha were distinguished while PP cells and undefined cells are mixed to some extent which is also consistent with original paper.

**Figure 3.**
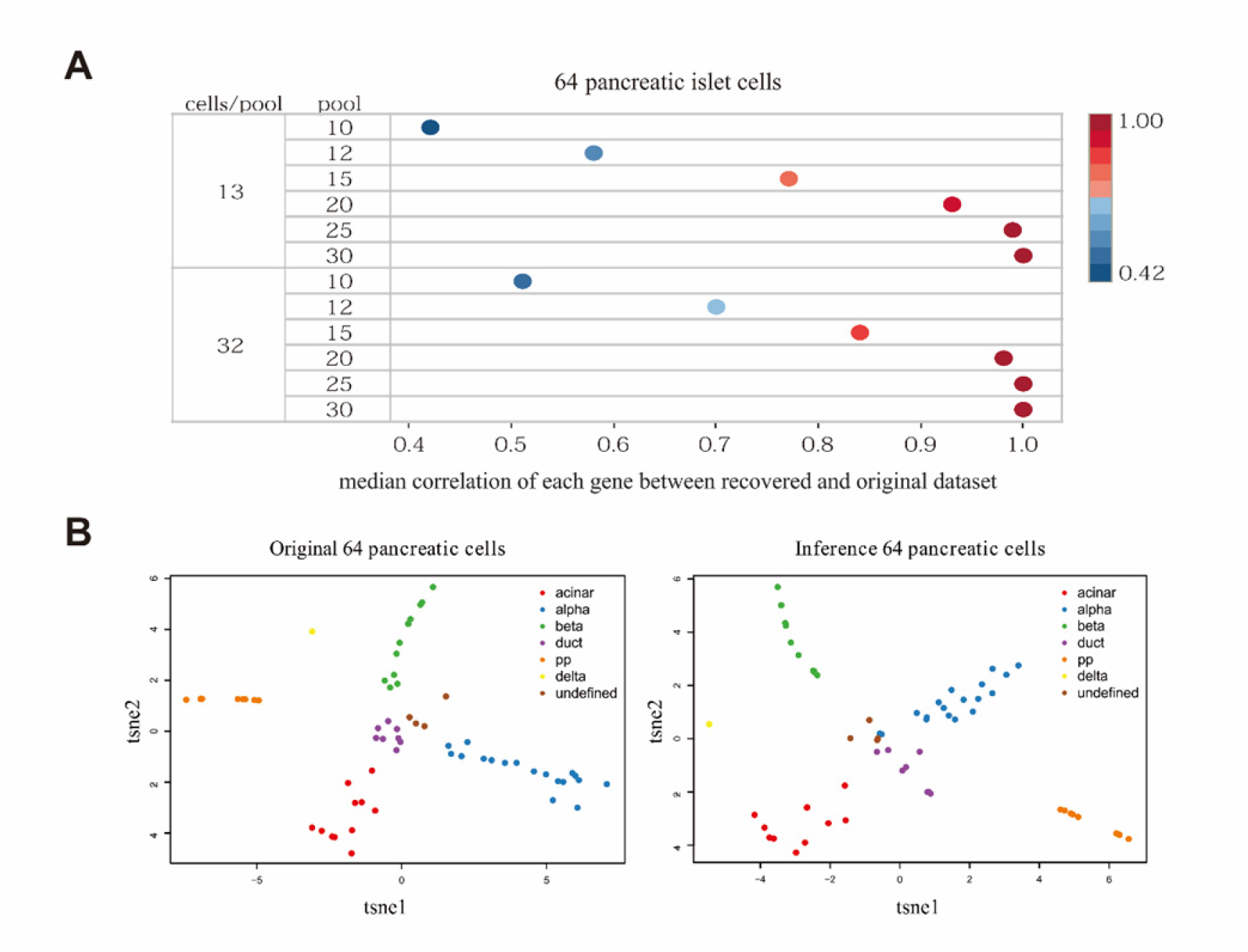
64 single cells toy example using compressed sensing method. (A) Pearson correlation for each gene between original and inferred SCEP with different pool numbers and cell numbers in each pool. (B) A low dimension embedding visualization for both original data and inference. 30 pools were used, 13 cells in each pool.

Overall, these results showed that by using only compressed sensing method people can successfully reconstruct SCEP with high accuracy when enough pools were used. Therefore, we achieved the goal to cut down the library cost at least twice. Though only half of the cost was cut down and more experimental efforts were paid, the merit of our approach will be amplified when dealing with much more cells. However, challenges are still remained like how to recover the non-sparse genes of SCEP and efficiency when computing an enormous scale of single cells. Next, we will extend this method to tackle both problems.

### A Parallel Comprehensive Model to Speed Up: Model Design

To cope with the high time complexity of accurate recovery in traditional compressed sensing (Chen et al., 1998; Dantzig, 1963), especially when the design matrix *M* is large (like thousands of cells in a bunch), we developed a parallel computing paradigm which reduces the computation time dramatically while allow experiment designers to choose the best combination configuration based on experimental condition.

The scheme is illustrated in Figure 4A. In principle, suppose we intend to impute the SCEP of *n* cells with *m* pools, the dimension of pool design matrix *M* is *m* × *n*. We do not build *M* as a whole at a time. A series of *M_i_*, *i* ∈ (1: *q*) were built instead, which line on the diagonal of the original matrix. For each sub-block 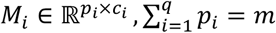, it is used as a pool design matrix for a subset of cells *c_i_*, which is non-overlap. In this manner, we bound all recovered *X_i_* by row to form the recover expression table as a whole.

**Figure 4.**
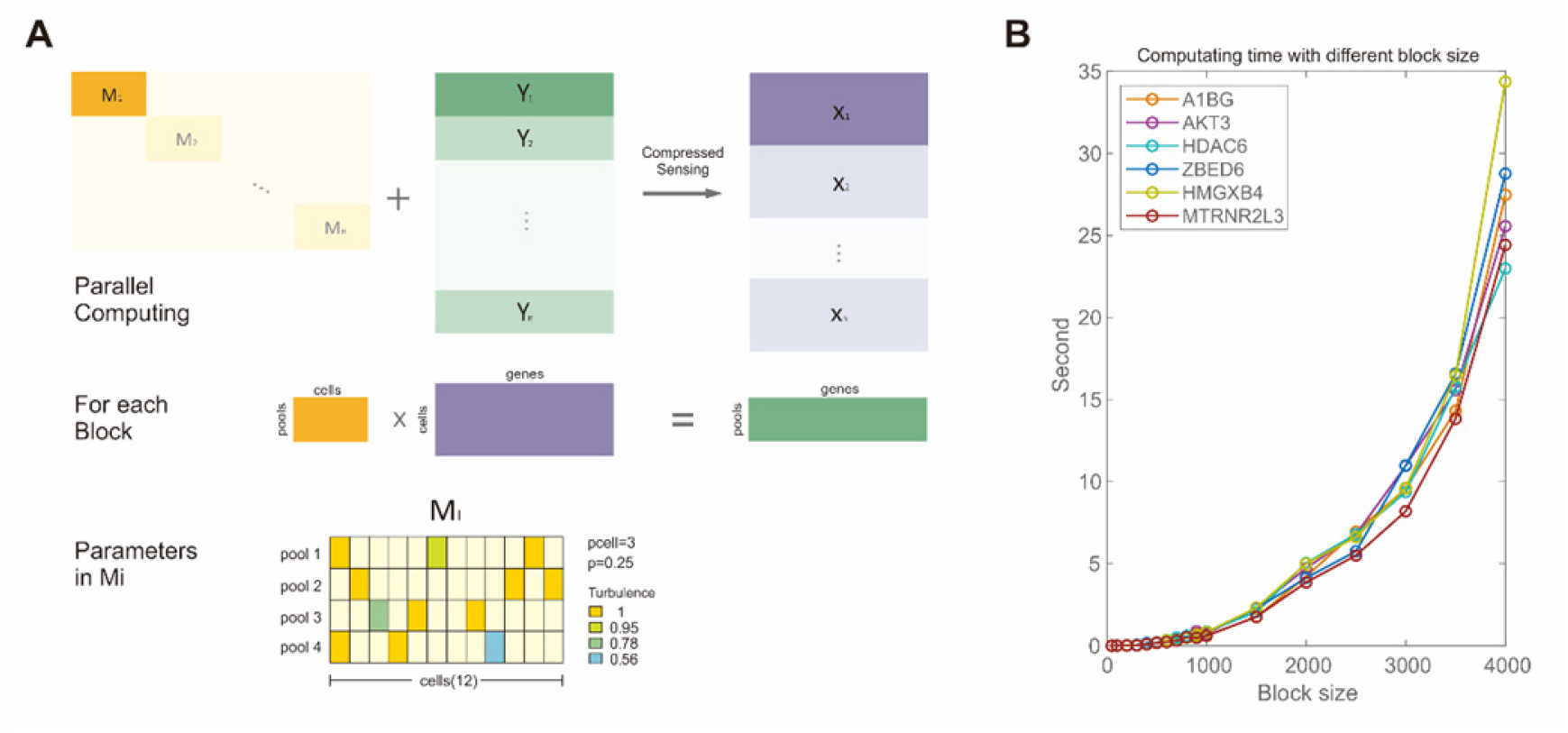
Parallel comprehensive model scheme lifts the efficiency. (A) Parallel computing scheme. A series of measurement matrix *M_i_* to observe a subset of cells. Using compressed sensing methods to recover each subset cells *X_i_* with *M_i_* and composite expression level of *Y_i_*, we stack all *X_i_* by rows to get an overall recovery of *X. pcell* indicates the number of cells contained in a pool, *p* stands for the percentage of *pcell* divide the number of all cells, which is 0.25=3/12 in this example. Different color stands for actual amount of lysate of each cell that is added in a pool, deviation from 1 is regard as noise in *M*. (B) Parallel model for speeding up. Six genes were randomly chosen to test effectiveness of parallel computing. For simplicity, we only chose first 4000 cells for simulation. Smaller block size indicates less cells in a group which also means more parallel threads. Computation environment is 8g RAM, Matlab 2018a.

This comprehensive model improved the flexibility of the experiment design. In order to determine the minimal pool size which is informative of original data, we set up few parameters. By the simulation using a 5063 single cells dataset (Zheng et al., 2017), we assessed how different parameters influence the performance. Four main parameters to concern (Figure 4A):

1. The number of different single cells in each pool: ***pcell***, which roughly equals to the number of non-zero value in each row of *M_i_*. Since we generated *M_i_* using a random scheme, ***pcell*** is an approximate value and may vary a little across different pools.
2. The sparsity level for each pool: ***p***. It determines the sub block size together with ***pcell*** by *size*(*columns of M_i_*) = *p/pcell*.
3. Total number of pools ***m***. This number is of critical importance and has huge impact on the final performance of our method. It is also the direct indicator of the library cost.
4. Sample add-in perturbation: ***t*.** It indicates the possible experimental error. It is a value between 0.5 to 1 and represents the variance and noise when samples were added into each pool. For example, *t* = 0.8 means the actual aliquot of cell is randomly drawn between 0.8 to 1.2.

For the number of cells in each pool *pcell*, it is a tradeoff between accuracy and experimental labor. Generally, a larger ***pcell*** value means each pool as an observer has more antennas to sense the signal (expression profile for one gene), which leads to a higher probability to maintain the structure and accurate value of original signal. However, to add more various type of cells in a pool may increase the efforts for experimental pooling.

Experimental noises are viewed as the noise of *M* matrix. The volume of a partition of one single cell is small and the RNA concentration and distribution in the partition might deviate from equal division. Here we evaluated the performance taking these noise into considerations.

### A Parallel Comprehensive Model to Enormous Data Set: Results

We focused on 5063 T cells isolated from different tissues and locations from liver cancer patients with deep depth scRNA-seq data as a showcase here (Zheng et al., 2017). Our comprehensive parallel model was applied on these cells with two goals same as the toy example: (1) Inferring exact value. (2) Finding a minimal possible number of pools that distinguishes different cells. Both goals were accessed by different combination of few parameters mentioned above. Finally, we also examined the running time under different block size to evaluate the speed up for parallel model.

#### Level One: The Accuracy Of Recovery

The results using a parallel scheme on 5063 single cells data set are shown in Figure 5A. To achieve the goal of reducing library consumption, we started the simulation from 2000 pools which is twice less than 5063 libraries as before. We chose this amount based on the sparsity level of *X* and the compressed sensing property which indicates that when pool *m* satisfies 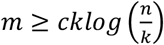, then people can recover the data of high accuracy with confidence (Supplementary method details). We then reduced the libraries gradually to assess the recovery performance. The reason we believe that use libraries less than the starting pool-number would still work is based on empirical knowledge of single cell profile density distribution as well as practical examination (Figure S1).

**Figure 5.**
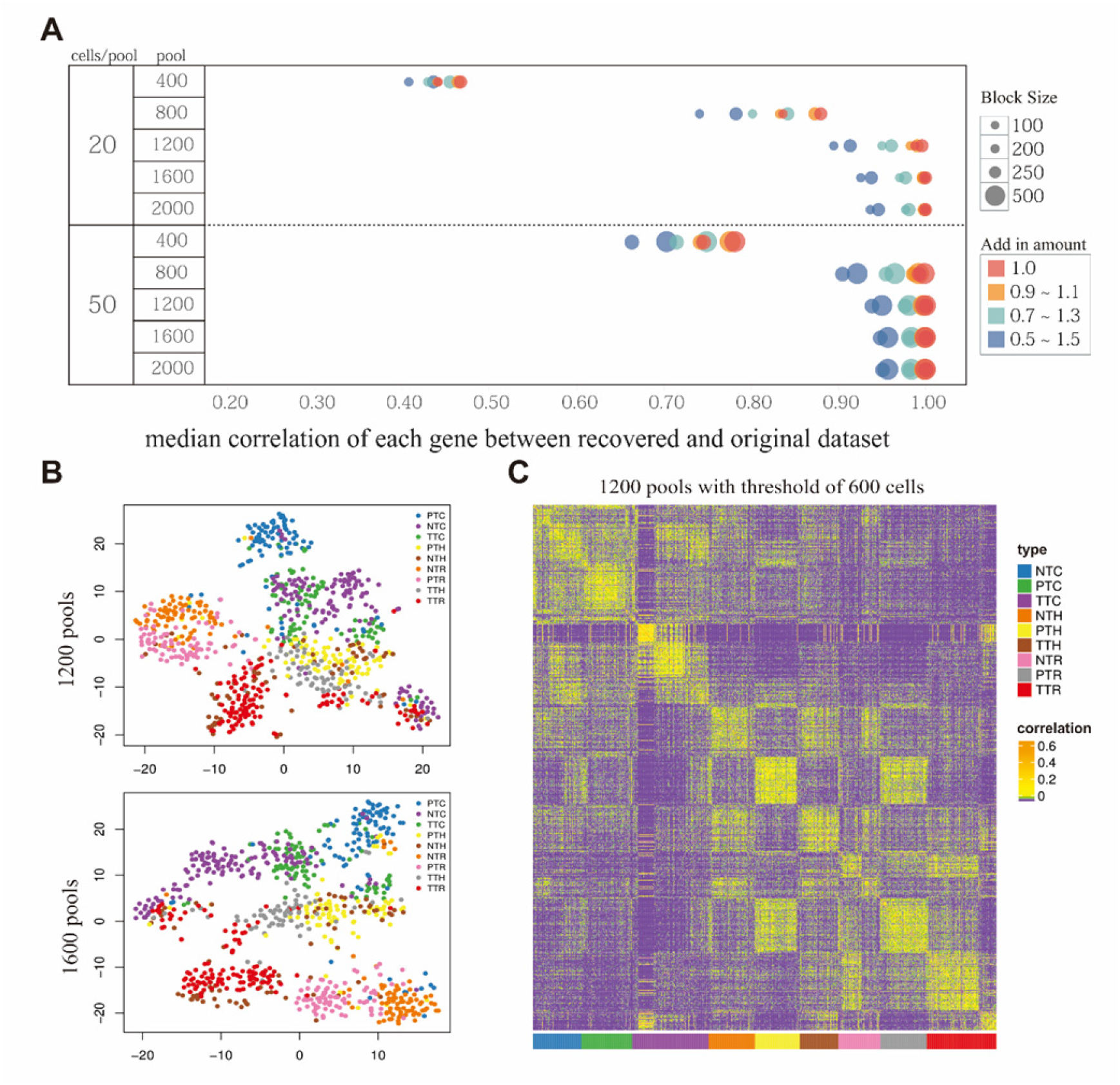
The results using a parallel scheme on 5063 single cells data set. (A) An Overall result for parallel comprehensive model. (B) Low dimension embedding visualization of recovered single cell expression profile using parallel comprehensive model of 1200 and 1600 pools. Each dot represents one single cell, tags were assigned according to original dataset indicates cell types and locations. (See also Figure S2). (C) Cell-cell correlation heatmap by inference with 1200 pools. Cell type patterns were annotated according to previous work. (See also Figure S4).

We can easily spot that the number of overall pools has a profoundly influence on the accuracy, and there was a sharp decrease from 800 to 400 pools.

As expected, *pcell* was also positively correlated with the accuracy. The more probes (cells in one pool) we used to sample a signal, the more information we got which also means a better performance for the recovery. We also investigated the add-in sample perturbation *t* ∈ (0.5,1) in experiment, the result showed that the accuracy was robust to the perturbation, median correlation can still achieve above 0.9 when the SNR decreased down to 2 (over 800 pools) (Figure 5A). In summary, the performance of our model lift with more pools, more cells in a pool, larger block size and less perturbation when add in sample.

#### Level Two: Rough Structure

One prominent task in single cell sequencing is to distinguish cell types according to their gene expression profile. It is no need to have the exact value to tackle this problem as long as we keep relative relationship of different genes. To be more specific, only genes that differently expressed across different cell types will play crucial role in cell type identification. Here we only kept part of genes for inspection with following reasons.

The first reason is to our knowledge that most of the housekeeping genes express at a relatively stable level across all kinds of cells, hence it is usually used to be a ruler for different cells under different conditions to rectify other genes’ expression level. This idea is used like spike-in strategy in qPCR, DESeq2 normalization pipeline (Love et al., 2014). These genes are usually not informative for classification. The second reason is that compressed sensing methods is originally designed to recover those sparse signals. From simulation we found that, the algorithm returned a sparse result even if the original signal was dense (a gene expressed cross a large number of cells) (Figure 6A). These distorted results will not reflect the real sparse level, so we evaluated the performance without these dense genes by setting different thresholds.

**Figure 6.**
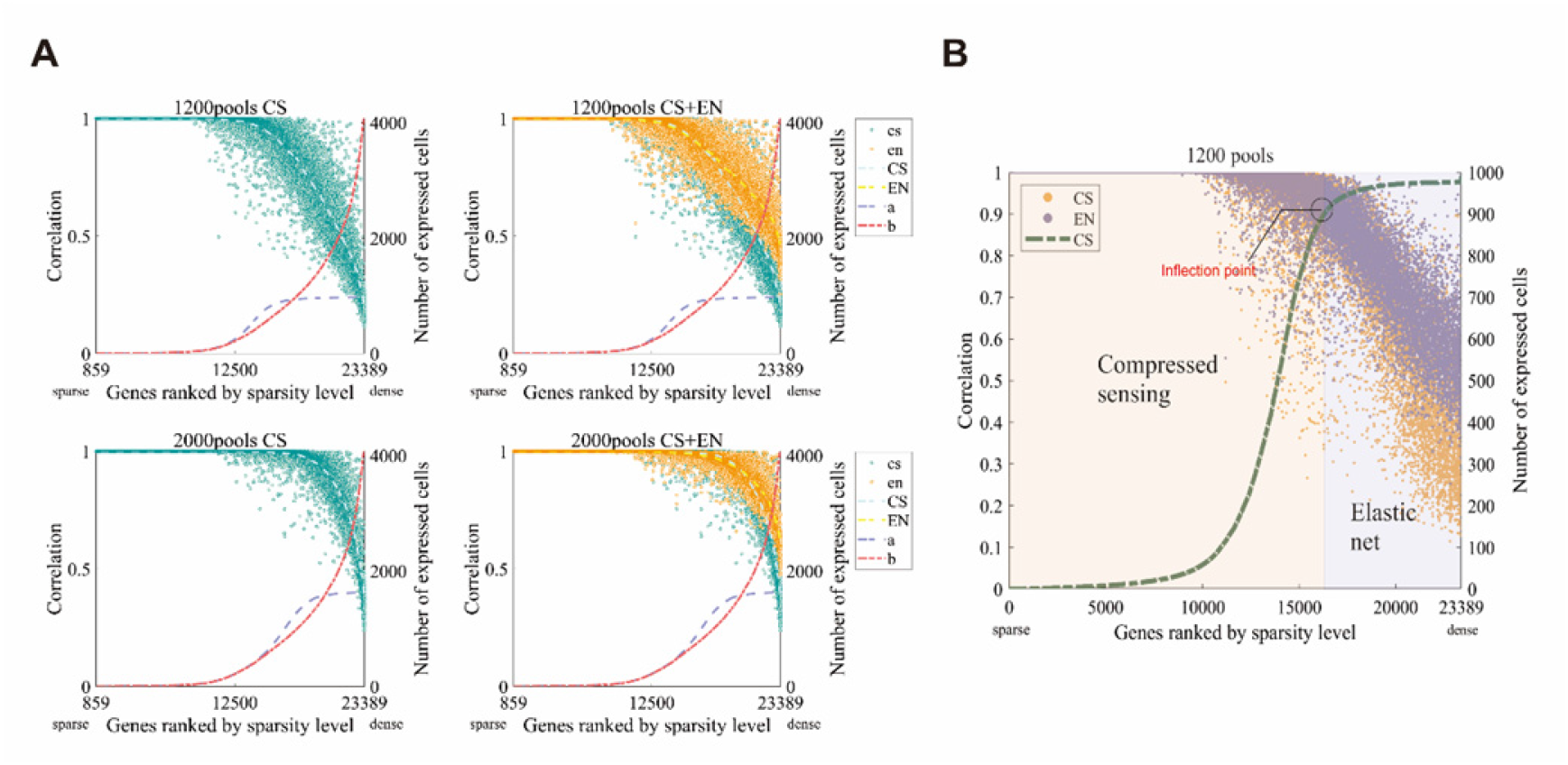
A combined model tackling both sparsity and density. (A) Comparison between compressed sensing only method (left column) and combined model (right column). Each dot represents a gene defined by it’s sparse level and recovery accuracy. Genes that not expressed in any cells (858 genes) and unqualified cells (as original paper) were excluded. Dash *b* line indicates the sparsity level in original data and the dash *a* line is for recovered data. (B) Combined model scheme. The dash line indicates expressed cells numbers of each gene from the results of Compressed sensing only method and ranked by sparsity level. Inflection point is estimated as a threshold to decide how to combine two results.

We chose the profile of patient 1116 (818 cells) for downstream analysis as the original paper did. Our model successfully distinguished nine different cell types by using only sparse genes (Figure 5B, Figure S2). The threshold to define sparsity is extended bilaterally from a half of number of pools we used in experiment. The pivot number is chosen based on theory but proved to work well in experiment. We show that using only 1600 libraries instead of 5063 libraries can distinguish each cell type, thus reduced two third of the cost. More interestingly, by using 1200 or less pools, even though we can’t get the distinct classification for each type, informative patterns were found. NTR and PTR (CD4^+^CD25^high^ cells) are always grouped together, and TTH, PTH and NTH (T helpers cells) also aggregate. Cell-cell correlation was also investigated (Figure 5C, Figure S4), even though the correlation between was pretty low, the intra-type correlation can still be discriminated obviously from inter-type cells (where the correlation is usually near zero).

#### Speeding Up

Six genes were picked out as an example to test the computation time using different block size in parallel scheme. As shown in Figure 4B, Time consuming increases as block size gets bigger, since block size is negatively correlated with parallel threads. Notice that the trend is non-linear, take gene HMGXB4 as an example. The computation time for recovering this gene speeds up near 7 times (from 5 to 35 seconds) when block size gets twice bigger (from 2000 cells per block to 4000).

So that using smaller blocks with more parallel computing threads can reduce time. However, people should notice that more computing resources were needed and less accuracy were achieved when using too many threads (Figure 5A).

In summary, by using parallel comprehensive model we achieved both high accuracy and maintained the core structure of the data whilst reduced the computing time significantly.

### One Stone Kills Two Birds: A Combined Model Tackling Both Sparsity and Density

Despite only some sparse genes are informative enough to cell type classification as discussed above, dense genes may also be desired for downstream analysis. Here we provided a new scheme which combined compressed sensing method for sparse genes and elastic net method for dense genes. Basically, we picked out those relatively dense genes in each sub-block from previous compressed sensing framework and recovered them again by elastic net.

The results for this adaptive method are shown in Figure 6A and Figure S3. We ranked every gene by their sparsity level from the sparsest on the left to densest on the right. Compressed sensing took advantage for sparse recovery so that both methods achieve same high accuracy for sparse genes (Figure 6A). However, a combined model outperformed pure CS method for dense genes recovery.

Highly variable genes were compared from original and recovery data by considering both variability and average expression level (Satija et al., 2015). Under the same threshold, increasing number of pools showed greater ability to capture same variable genes as original data (Figure S5) which may have huge impact on downstream analysis.

In summary, a combined model is more adaptive and took advantage of both CS and elastic net methods at the same time so that achieves a higher accuracy. This model is also more flexible that people are allowed to set different threshold for partition between sparsity and density, balance between lasso and ridge regression.

### A Laborious Model for Higher Accuracy

As we mentioned above, a necessary and sufficient condition for recovering *x* with high probability requires measurement matrix *M* satisfies RIP *(restricted isometry property)* condition (Baraniuk et al., 2008; Candes et al., 2006). A randomly generated Gaussian or Bernoulli matrix meet this requirement. A random Gaussian matrix *M*’ is usually full but with a better ability to capture more information from *Y*. Yet, from experimental perspective, a full pool design matrix *M* requires more intensive work for adding sample in each pool.

Here we still provide a full pool design matrix scheme for possible usage. We compared the results (Figure 7) using 400 pools and 800 pools which performed ordinary when using Bernoulli matrix. New *M’* with original *Y* outperformed previous scheme in every aspect, especially when the block size was small (like 200 columns of each M). This can be interpreted as just a proportion of detectors in a small M cannot even make sure that each cell was detected at least once. However, a random full matrix can always guarantee that.

**Figure 7.**
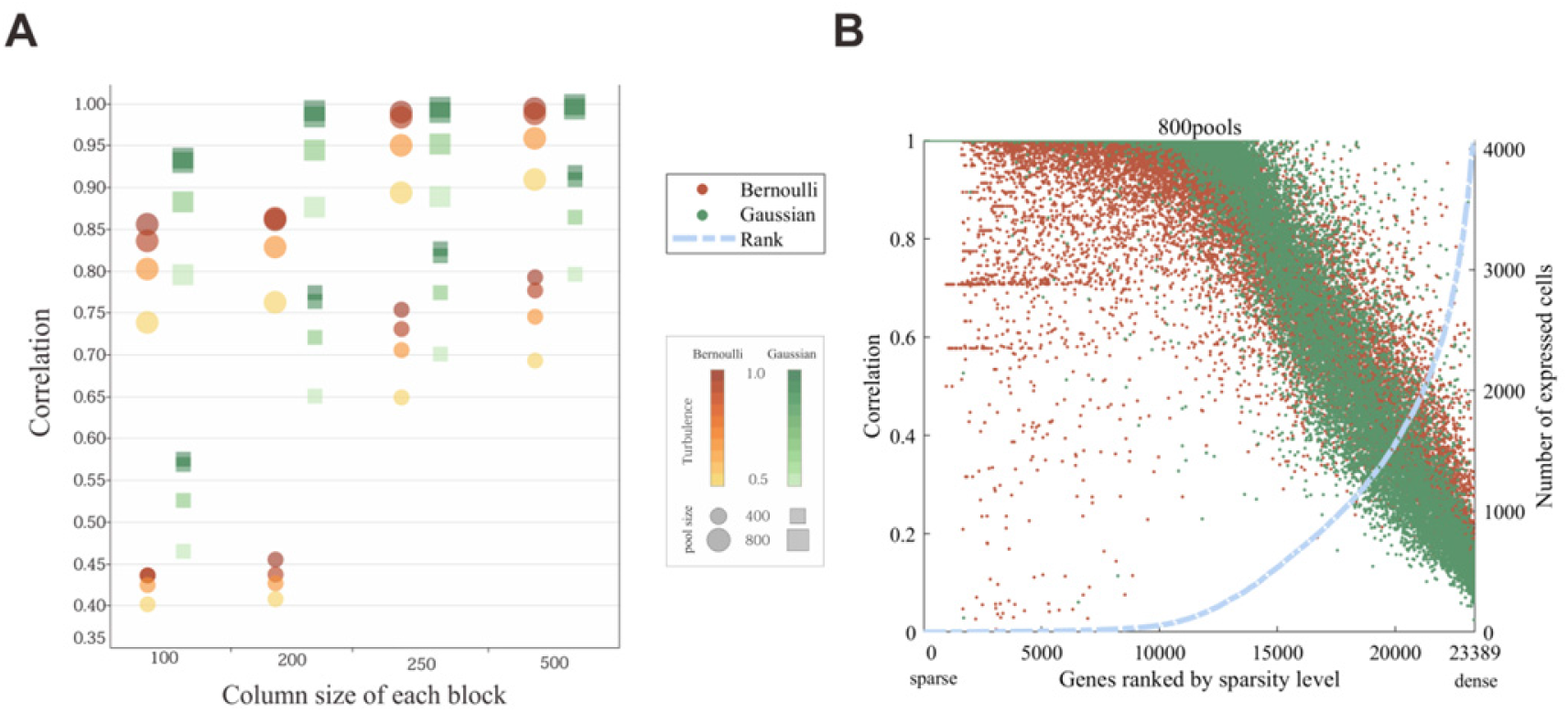
Gaussian design matrix model improved the accuracy. (A) Results comparison between using random binary pool design matrix (Bernoulli) and full matrix with 4 concentrations randomly distributed to approximate Gaussian distribution (Gaussian). (B) Correlation analysis of recovery accuracy with two design matrix schemes with 800 pools.

Even though a full matrix *M’* can achieve a better performance, there is a tradeoff between the experiment efforts and inferring accuracy. One single cell is represented using aliquots of different volumes in all pools. Alternatively, with the increasing demand for cell’s nucleic acid in a full matrix, amplifications can be applied before pooling, which might introduce more variance and bias.

## DISCUSSION

To close the gap between cost and efficiency of gene detection in single cell sequencing, we adopted the grouping-based pooling strategy for SCEP inference. Only less than a half of sequencing libraries were needed while the gist information captured. Unlike massively parallel approaches like Drop-Seq which could only detect limited kinds of genes, this pooling strategy gets a more extensive expression profile. Meanwhile, the maintainability of full length RNAs improves the accuracy of gene expression profiles in comparison with the massive approaches which could only get short sequences adjacent to the barcodes. In this grouping-based pooling strategy, the number of pools is much less than the number of cells, but increases slightly along with the increase of cells (Kang et al., 2018; Poli et al., 2009; Satija and Shalek, 2014). Therefore, if the number of cells is extremely huge, the cost used by pooling strategy should still be taken into account.

On the basis of compressed sensing framework, the inference process can be transformed to solve an ill-conditioned linear equation: *Y* = *MX*. The only unknown coefficient matrix *M* is a connection between cells and pools, it is an abstraction of this measurement transformation. The designed Bernoulli matrix is a binary matrix which resembles a switch between choosing a single cell to a pool or not. The designation also adopts the idea of overlapping from group testing methods. Compared with a binary matrix, a random Gaussian matrix achieves higher accuracy but requires more laborious operations in practical experiments. There is always a tradeoff between numeric accuracy and experimental efforts. The same is true when we want a better performance by increasing the number of pools and add in more cells in each pool. The automatic sampling systems might relieve this dilemma between performance and experimental effort in the future.

To solve this under-determined system, CS is used for sparse genes recovery, and elastic net is aimed for non-sparse genes. With a progressive approximation algorithm, CS is efficient considering time complexity, and these inferred sparse genes are sufficient to maintain the core structure of original data independently which has been proved by our investigation. The non-sparse genes are also useful for clustering and that are crucial for downstream analysis. Therefore, it is worth improving the inferring accuracy of non-sparse genes through elastic net. Of course, the dense genes can’t be fully recovered on the basis of information theory without any assumptions.

Extraction features from datasets for cell type identification has been a relish challenge in cell analysis. Previous studies have suggested that highly expressed and highly varied genes contain most crucial information for classification (Satija et al., 2015). Although it is a plight to distinguish those highly varied genes from single cell expression profiles which are usually sparse due to a high percentage of drop-out events, our simulation demonstrated that sparse genes were also capable to capture the gist of structure. The results suggested that the highly varied genes and the sparse genes might be greatly overlapped, and the sparse property is a potential index to extract the genes which contain most crucial features for classification.

Cells from same individuals, tissues or positions, or with similar phenotypes revealed similar expression profiles in previous studies (Li et al., 2016; Zheng et al., 2017). More resemblance in gene expression is seemed to be carried by cells with similarity. On the other hand, genes are tended to gather in groups in expression profiles which suggests that genes in a group are strongly correlated. Ideas were raised by leveraging these dependencies like eigengenes (Steve and Peter, 2007) and gene modules (Cleary et al., 2017). Both the similarity in cells and the correlation between specific genes can be utilized as pre-information for low rank compressed sensing recovery (Fazel et al., 2008), which might lead a better performance for dense genes recovery.

To sum up, single cell expression profile analysis is an emerging and exciting area to explore with its intrinsic features like sparsity. Traditional analysis methods should be improved and adapted based on this property. Our work offered a comprehensive scheme on recovering an expression profile composed of both sparse genes and non-sparse genes with a dramatic decrease on library consumption. This frame work can also be widely adapted to other single cell atlas including proteomics, lipidomics and metabolomics as long as a sparse feature is presented. More works in this area are enthusiastically to be seen for the coming years.

## STAR METHODS

Detailed methods are provided in the online version of this paper and including the following:

- **KEY RESOURCES TABLE**
- **METHODS DETAILS**

○ Dictionary Design
○ Toy example simulation processing
○ Parallel comprehensive model on large dataset
○ Combined model processing
- **QUANTIFICATION AND STATISTICAL ANALYSIS**

○ Sparse gene set identification
○ Single cell expression profile normalization
○ Differential expressed gene sets identification
- **DATA AND SOFTWARE AVAILABILITY**

○ Software Availability

## SUPPLEMENTAL INFORMATION

Supplemental Information includes five figures and can be found with this article online.

## AUTHOR CONTRIBUTIONS

X.W. developed the algorithms, performed the analysis, wrote the manuscript. W.X. developed the algorithms. J.T. conceived of the project, designed the analysis and revised the manuscript. X.S. conceived of the project. J.T. and Z.L. guided research.

## ACKNOWLEDGEMENT

This work was supported by National Key R&D Program of China (No. 2016YFA0501600), National Natural Science Foundation of China (No. 61571121), Key Research & Development Program of Jiangsu Province (No. BE2016002-3), and Fundamental Research Funds for the Central Universities of China. We thank Prof. Dehan Kong for suggestion for further discussion on low rank recovery, Dr. Jiapeng Chen for helpful discussions, Dr. Jianhua Yin for help with graphic, and An Ju and Charlotte Chen for language polishing.

## STAR METHODS

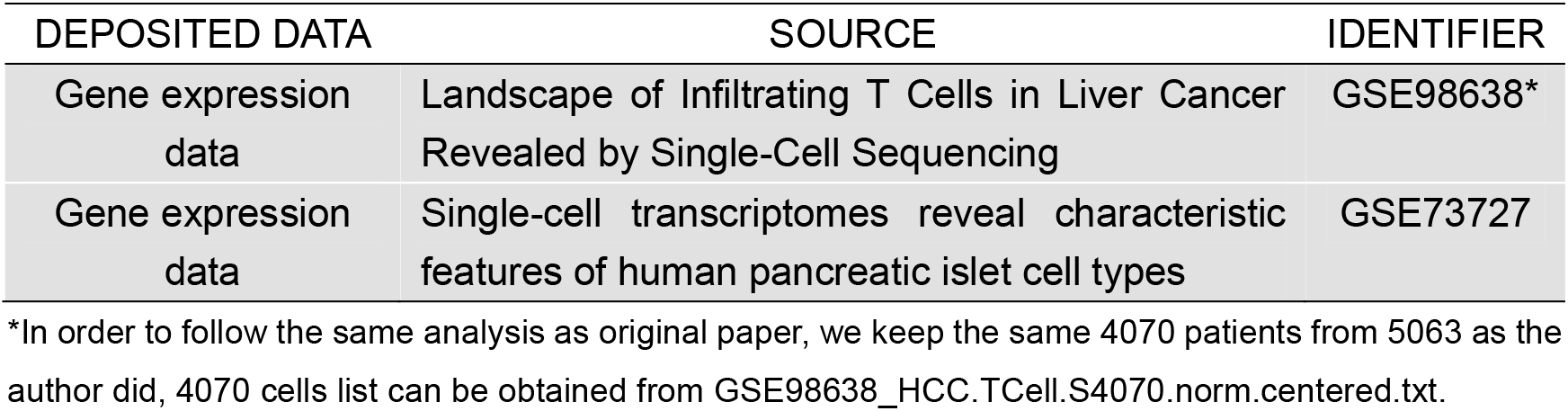
KEY RESOURCES TABLE

KEY RESOURCES TABLE
| DEPOSITED DATA | SOURCE | IDENTIFIER |
| --- | --- | --- |
| Gene expression data | Landscape of Infiltrating T Cells in Liver Cancer Revealed by Single-Cell Sequencing | GSE98638* |
| Gene expression data | Single-cell transcriptomes reveal characteristic features of human pancreatic islet cell types | GSE73727 |

\*In order to follow the same analysis as original paper, we keep the same 4070 patients from 5063 as the author did, 4070 cells list can be obtained from GSE98638\_HCC.TCell.S4070.norm.centered.txt.

## METHOD DETAILS

The inference procedure is designed to recover a single cell expression profile by a known design matrix and a measurement matrix for all pools.

### Dictionary Design

Toy example is generated based on a 64 single cells dataset (GSE73727). We built *M* by first generated a gaussian random matrix with dimensions of *pools* × *cells*, then set a threshold to transform gaussian matrix to binary matrix by value comparison which return a logic value. In parallel comprehensive model, for each sub *M_i_*, we used the same method as above. *p* is set between 0 and 1 to determine the sparsity level of each pool. *p* = 0.1 represents around 10% cells were picked in each pool. In full pool design matrix, we first generate a vector of equal length of cell numbers in each sub block. Each vector contains near equal number of {1, 0.1, 0.01, 0.001}. Then we constructed *M* whose rows are random permutations of the vector.

### Toy Example Simulation Processing

A transcript expression level table for 64 single cells of different pancreatic islet cell types (Li et al., 2016) dataset EV2 was downloaded. We then collapsed the dataset based on the gene symbol “ensG” by *plyr* package in R. Using this computed cell-gene expression table *X*, and multiplying by designed measurement matrix *M*, we then built up composite measurement by *Y* = *M* × *X*. Mwas varied on the number of pools from 12 to 30, and number of cells in each pool was 13 or 32. Basically, the way we decided how to allocate different cells in each pool is by simulating numbers from a uniform distribution *u* ∈ [0,1] and then set a threshold. For all positions, if the value is less than the threshold, it would be set to one otherwise to zero. The threshold *p* can be set between 0 to 1, and the number of cells in each pool equals *p* times number of all cells. Then 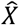 is solved by CS method. (https://github.com/Irenexzwen/SigRec)

### Parallel Comprehensive Model on Large Dataset

We collected data of 5063 single cells from NCBI (GSE98638) (Zheng et al., 2017), and then excluded unqualified single cells followed the original paper for downstream analysis. To implement a parallel model, we first constructed *M_i_* based on four parameters introduced before. To implement parallel scheme, we partitioned original expression profile by rows into sub groups of cells’ expression profile *X_i_*, whose row number equals the column number of *M_i_*. Each block was then computed using same scheme as toy example. Finally, we bounded all recovered 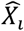 by rows to form the SCEP. For the turbulence *t* ∈ (0,1), we replace value 1 with a random number sampled from a uniform distribution *u* ∈(*t*, 1 – *t*).For classification visualization, we chose patient 1116 followed the original paper. The sparsity distribution of all genes were calculated once we got the recovered SCEP and set expressed cells cut off according to 1^st^ and 3^rd^ quantile value.

### Combined Model Processing

Elastic net regression was applied based on the results of purely compressed sensing. For each block of recovered 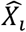, we counted the number of non-zero items of each gene across all cells and picked out those dense genes whose expression profile in Y are all non-zero. These selected genes were then regressed using elastic net by tuning the hyperparameters λ and α using cross validation. After regression, we stuck them back to the original position in each sub regressed 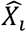 and then bounded all 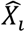 by rows together to get the recovered 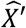. Next, we examined the result of compressed sensing only method to find a inflection point as a threshold. Genes whose expressed cells numbers beyond this threshold were viewed as “inappropriately regressed” so that we substituted these genes expression inference with the results of elastic net regression to get a final result 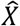 (Figure 6B).

### Full Pool Design Matrix Scheme

In full pool design matrix, first row in new *M*’ matrix was generated by randomly sampling from {1,0.1,0.01,0.001} with equal frequency, which represents four concentration gradients. Then we constructed *M_i_* with all rows are random permutation of the first row. By this manner, we created a near random matrix but only with four possible value for experimental convenience. Following procedures were exactly the same as comprehensive parallel model with know *Y*.

## QUANTIFICATION AND STATISTICAL ANALYSIS

### Sparse gene set identification

After we got recovered single cell expression profile from compressed sensing method, for each gene *i*, we first calculated the number of cells that expressed this gene (non-zero items in each gene) as *c_i_*, then sorted all genes by *c_i_* increasingly to get a vector C_i_. Function Q is defined as the quantile function. For specific pool numbers *m*, Q_C_*i*__.(0.5) was chosen as a pivot threshold(median) and extended the value towards Q_C_*i*__.(0.25) and Q_C_*i*__.(0.75) by equal 5 intervals. All genes that has a *c_i_* < threshold will be preserved for downstream analysis like t-SNE and correlation analysis.

### Single cell expression profile normalization

Zero-inflation property of single-cell expression profile (high proportion of zero counts due to biological or technical reasons) account for a poor performance when using bulk-seq normalization methods (Vallejos et al., 2017). However, to follow the same strategy as original research using DESeq2 (Anders and Huber, 2010), we tailored the method to meet our sparse recovered results. Briefly speaking, DESeq2 calculates global scale factors based on a pseudo-reference which is composed of geometric mean of specific genes over all samples, unfortunately, it’s too rigorous for a zero-inflated SCEP to meet this requirement. We circumvent this dilemma by choosing the densest cell from our inference (expressed most genes) as reference cell. Then size factor for each cell is the median of gene expression level ratio (only genes that expressed in both cells) between two cells. After that expression level for each gene of a cell is modified by multiply the size factor calculated above.

### Differential expressed gene set identification

We identified highly variable genes followed the same method as Seurat. Basically, both average expression and dispersion of each gene are taken into consideration. Parameters were chosen as default value from the toolkit (x.low.cutoff = 0.0125, x.high.cutoff = 3, y.cutoff = 0.5). We tested the combined model results (4070 cells and 23389 genes) by this method generated Figure S4.

## DATAT AND SOFTWARE AVAILABILITY

### Software availability

Analysis and code scripts can be found at https://github.com/Irenexzwen/SigRec.

